# Identification of two genomic cryptotypes of *Plasmodium malariae* in Africa

**DOI:** 10.64898/2026.03.23.713578

**Authors:** Margaux J. M. Lefebvre, Céline Arnathau, Sandrine Houzé, Benoit de Thoisy, Camila González, Silvia Rondón, Andrés Link, Arnab Pain, Michael C. Fontaine, Franck Prugnolle, Virginie Rougeron

**Affiliations:** MiVEGEC, Univ. Montpellier, CNRS, IRD, Montpellier, France; Department of Archaeogenetics, Max Planck Institute for Evolutionary Anthropology, Leipzig, Germany; Université de Paris, MERIT, IRD, Paris, France, AP-HP, Centre National de Référence sur le paludisme, hôpital Bichat-Claude-Bernard, Paris, France; Institut Pasteur de la Guyane, Laboratoire des Interactions Virus Hôtes, Cayenne, Guyane, France; Centro de Investigaciones en Microbiología y Parasitología Tropical (CIMPAT), Universidad de los Andes, Bogotá D. C., Colombia; Laboratorio de Ecología de Bosques Tropicales y Primatología (LEBTYP), Universidad de los Andes, Bogotá D. C., Colombia; Pathogen Genomics Laboratory, Biological and Environmental Sciences and Engineering, King Abdullah University of Science and Technology, Jeddah Makkah, Saudi Arabia; International Institute for Zoonosis Control, Hokkaido University, Sapporo, Japan; Groningen Institute for Evolutionary Life Sciences (GELIFES), University of Groningen, Groningen, The Netherlands; REHABS, International Research Laboratory, CNRS-NMU, George Campus, Nelson Mandela University, George, South Africa; Sustainability Research Unit, George Campus, Nelson Mandela University, George campus, Madiba drive, 6529 George, South Africa

**Keywords:** *Plasmodium malariae*, *Plasmodium brasilianum*, population genetic, malaria, Africa, adaptation

## Abstract

*Plasmodium malariae* is a neglected human malaria parasite that causes persistent, often asymptomatic infections and remains difficult to diagnose. Despite being generally associated with lower prevalence and severity than other malaria parasites, *P. malariae* represents a significant public health concern, particularly in Africa, but also as a zoonosis in South America with monkey-adapted *Plasmodium brasilianum. Plasmodium malariae* and *P. brasilianum* population genetic structure, evolutionary history, and adaptive potential remain poorly understood, largely due to the historical scarcity of whole-genome data. By screening 226 monkey samples from two Latin American countries, we identified 20 *Plasmodium*-positives across multiple primate species, highlighting the persistence of this parasite in sylvatic transmission cycles. We also investigated the evolutionary history and genetic diversity of *P. malariae* using whole-genome sequencing data. By combining 79 newly sequenced genomes with 248 publicly available genomes, we analyzed a filtered dataset comprising 179 *P. malariae*, two *P. brasilianum*, and two *P. malariae*-like genomes. Population structure analyses revealed the presence of two genetically distinct but recombining clusters across African *P. malariae* populations. These clusters occur across multiple African countries at varying frequencies, without clear geographic segregation. Genome-wide scans of genetic differentiation and selection further identified numerous cluster-specific signatures of adaptation, including loci putatively involved in interactions with human hosts and mosquito vectors. Our results provide the first evidence for fine-scale population substructure within African *P. malariae* and reveal ongoing adaptive processes that may contribute to its persistence and transmission. By uncovering previously unrecognized genetic diversity and selection patterns, this study highlights the importance of population genomic approaches for understanding the evolutionary dynamics of this neglected malaria parasite.

**Author summary:** Human malaria is most often associated with few well-studied parasites (*Plasmodium falciparum* and *Plasmodium vivax*), while other species have received far less attention. One such species is *Plasmodium malariae*, which often causes long-lasting infections with few or no symptoms. Because infections are difficult to detect, the true impact of this parasite is likely underestimated, particularly in Africa, where it is widespread. A closely related form, known as *Plasmodium brasilianum*, infects monkeys in South America with transmission between humans and wild animals. In our study, we analyzed the parasite genomes from infected people and monkeys to better understand the genetic diversity of these parasites, their population structure, and the molecular evidence of adaptation. We found evidence that *P. brasilianum* continues to circulate in several wild monkey species in South America. By analyzing nearly 300 genomes, we discovered that African *P. malariae* consists of two genetically distinct but recombining groups that coexist across the continent. These groups differ in specific regions of their genomes, including genes likely involved in interactions with human hosts and mosquito vectors. Our results reveal cryptic genetic diversity and structure with evidence of ongoing adaptation. These findings highlight the importance of including neglected parasites in malaria surveillance and control efforts.

## Introduction

With 272 million cases estimated and approximately 500 000 deaths reported in 2024 [1], malaria remains one of the most devastating infectious diseases worldwide. Although it is not generally classified as a neglected tropical disease, the overwhelming majority of cases and fatalities are attributable to two species, *Plasmodium falciparum* and *Plasmodium vivax* [1]. In contrast, other human-infecting malaria parasites, including *Plasmodium malariae, Plasmodium ovale curtisi* and *Plasmodium ovale wallikeri* [2], remain comparatively understudied resulting in substantial gaps in our understanding of their epidemiology, evolutionary dynamics, and adaptive potential. Among these neglected parasites, *P. malariae* is particularly enigmatic. Infections are predominantly asymptomatic [3,4], characterized by low parasitemia and can persist chronically for years [5]. Although *P. malariae* infections are often considered clinically mild, they can result in clinically significant outcomes, including anemia [6], nephrotic syndrome [5], and, in rare cases, mortality in children [6]. However, its true burden is likely underestimated due to diagnostic limitations and frequent misidentification [3]. Geographically, *P. malariae* is predominantly distributed in sub-Saharan Africa, where in several countries it is the second most prevalent malaria species after *P. falciparum* [7–10]. Beyond its distribution in humans across Africa, as well as in Asia, and South America, *P. malariae* is closely related to parasites infecting South American non-human primates (NHPs). These infections in American NHPs are traditionally referred to as *Plasmodium brasilianum* [7–9]. However, several molecular and genetic studies suggest that *P. malariae* and *P. brasilianum* may represent a single species complex [10–12]. Documented cross-species transmission between humans and NHPs further support this hypothesis [12], raising questions about zoonotic reservoirs, parasite persistence, and malaria control efforts in South America [12,13].

Despite its epidemiological importance, the population genetic structure, evolutionary history and adaptation of *P. malariae* remain poorly resolved. This knowledge gap largely reflects the historical scarcity of whole-genome data, limited geographic sampling across the parasite global distribution, and the technical challenges associated with generating genomic data from low-parasitemia infections. In addition to receiving comparatively limited research focus, *P. malariae* has not benefited from large-scale sequencing initiatives comparable to those conducted for *P. falciparum* and *P. vivax* [25,26]. Consequently, until recently, fewer than 25 *P. malariae* genomes were publicly available worldwide [21–24]. Although Ibrahim *et al*. [27] substantially expanded genomic resources by adding 222 *P. malariae* genomes, important limitations remain. While this effort increased representation across Africa, sampling within individual countries was variable and often limited, reducing the power to resolve intra-continental population structure. This situation is even more pronounced for *P. brasilianum*, for which genomic resources are extremely scarce, with only a single complete genome published to date [14]. This limited representation reflects substantial logistical and technical constraints, including restricted access to NHP blood samples, ethical and conservation regulations governing wildlife research, low parasitemia levels in natural infections, and high host DNA contamination. Moreover, because *P. brasilianum* has long been considered genetically indistinguishable from *P. malariae*, it has historically received limited priority in large-scale genomic initiatives. For all these reasons, a comprehensive dataset of both *P. malariae* and *P. brasilianum* is therefore essential for understanding their population genetic structure and host adaptations. This information is also necessary for developing effective malaria surveillance and control strategies.

In this study, we aimed to strengthen and densify African whole-genome representation of *P. malariae* (the region where human infections are most prevalent and where population structure remains insufficiently resolved) while simultaneously trying to generate additional genomic data for *P. brasilianum* to better characterize their genetic relationships with *P. malariae*. To do so, we screened 212 NHP samples and identified 20 PCR-positive *P. brasilianum* infections. In total, we generated 59 new *P. malariae* and 20 *P. brasilianum* whole-genome sequences. After integration with previously published datasets and final quality control, the resulting dataset comprised 179 *P. malariae*, two *P. brasilianum*, and two *P. malariae-like* genomes. Even if the screening of South American NHPs revealed substantial circulation of *P. brasilianum*, providing important epidemiological insights into its prevalence and host distribution across multiple primate taxa, only two high-quality *P. brasilianum* genomes were recovered, precluding detailed population genomic analyses. Regarding *P. malariae*, results uncovered two previously unrecognized recombinant genetic clusters within African *P. malariae* populations, consistent with cryptic lineage diversification across the continent. These clusters differ in genome-wide diversity, exhibit localized genomic differentiation, and show signatures of lineage-specific adaptation. These results highlight that African *P. malariae* populations, as African *P. falciparum* ones [15], are structured into distinct evolutionary lineages, revealing previously unnoticed genetic diversification within this neglected malaria parasite.

## Materials and methods

### Origin of samples, species identification, and ethical statements

For the 59 *P. malariae* samples, no specific consent was required because, in coordination with the Santé Publique France organization for malaria care and surveillance, the human clinical, epidemiological, and biological data were collected in the French Reference National Center for Malaria (CNRP) database and analyzed in accordance with the public health mission of all French National Reference Centers. The study of the biological samples obtained in the context of medical care was considered as non-interventional research (article L1221-1.1 of the French public health code) and only required the patient’s non-opposition during sampling (article L1211-2 of the French public health code). Genomic DNA was extracted using the Qiagen DNeasy Blood and Tissue Kit according to the manufacturer’s recommendations.

Regarding non-human primates (NHPs) samples, 212 were collected in French Guiana Those samples are registered in the collection JAGUARS (https://kwata.net/gestion-collection-biologique/, CITES reference: FR973A) managed by the Kwata NGO (accredited by the French Ministry of the Environment and the Prefecture of French Guiana, Agreement R03-2022-12-30-0007 and R03-2024-11-07-00036), hosted at the Institut Pasteur de la Guyane, supported by Prefecture de la Région Guyane and Collectivité Territoriale de la Guyane, and validated by the French Guianan prefectoral decree n°2012/110. Regarding samples from Colombia, ethical approvals for the collection of eight fecal and six blood samples were obtained by Universidad de los Andes, the National Environmental Licensing Authority of Colombia (ANLA) and the Centers for Disease Control and Prevention (permits numbers: 2017025578-1-000, 2017043863-1-000, 2017065795-1-000, 2017013727-1-000, 2017052943-1-000, 2017081458-1-000, 2017108650-1-000).

These NHP samples were predominantly from *Alouatta macconnelli* (n = 140) and *Saguinus midas* (n = 58), with lower representation of *Ateles paniscus* (n = 7), *Ateles hybridus* (n = 5), *Alouatta seniculus* (n = 5), *Sapajus apella* (n = 4), *Cebus versicolor* (n = 3), *Saimiri sciureus* (n = 2), *Aotus griseimembra* (n = 1), and *Pithecia pithecia* (n = 1). For the 226 samples from French Guiana and Colombia, genomic DNA was extracted using the Qiagen DNeasy Blood and Tissue Kit according to the manufacturer’s recommendations. *Plasmodium brasilianum* samples were identified by amplification of *Plasmodium cytochrome b* using nested PCR, as described in Prugnolle *et al*. [16]. The reaction products were visualized on 1.5% agarose gels stained with EZ-vision and sent for Sanger sequencing to confirm *Plasmodium* species (Eurofin MWG). This allowed the identification of 20 positive *P. brasilianum* samples.

### Selective whole genome amplification (sWGA), library preparation and sequencing

For the 59 *P. malariae* DNA samples, selective whole-genome amplification (sWGA) was performed to enrich parasite DNA from samples with low parasitemia using the *P. malariae*-specific protocol developed by Ben-Rached *et al*. [17]. This technique preferentially amplifies the *P. malariae* genome from mixed DNA samples while reducing host DNA contamination. Given the close genetic relationship between *P. malariae* and *P. brasilianum*, the same approach was also applied to *P. brasilianum* samples. The primer set consisted of ten 8–10 bp primers (PM1–PM10), each containing phosphorothioate bonds at the 3′ end to prevent exonuclease degradation. Each primer was initially prepared at 100 µM and combined into a Primer Set 1 master mix to achieve a final concentration of 1.225 µM per primer in a 50 µL reaction. For each reaction (50 µL total volume), approximately 50 ng of genomic DNA was used as input. The reaction mixture contained 17.5 µL of Primer Set 1 master mix, 5 µL of 10× phi29 enzyme buffer (New England Biolabs), 3 µL of phi29 DNA polymerase (30 U; NEB), 2 µL of 25 mM dNTP mix (ThermoFisher), nuclease-free water, and elution buffer (EB) adjusted to reach the final volume of 50 µL. Amplification was carried out using a “ramp-down” thermocycling program: the temperature was decreased stepwise from 35°C to 30°C (10 minutes per degree), followed by a 16-hour incubation at 30°C. Enzyme inactivation was performed at 65°C for 10 minutes, and reactions were held at 4°C until further processing. For each sample, the products of the two amplifications (one per primer set) were purified with AMPure XP beads (Beckman Coulter) at a 1:1 ratio according to the manufacturer’s recommendations and pooled at equimolar concentrations. Each sWGA library was prepared using the two pooled amplification products and the Nextera XT DNA kit (Illumina), following the manufacturer’s protocol. Then, samples were pooled, clustered, and sequenced on one lane of a Illumina Novaseq-6000 S4 with 2 × 150-bp paired-end reads (MGX Montpellier).

### Short-read mapping, SNP calling, and data compilation

We generated whole genome-sequencing data for 59 sequenced *P. malariae* isolates and 20 newly sequenced *P. brasilianum* samples. These were added to a compilation of previously published *P. malariae* genomic datasets: 222 samples from Ibrahim *et al*. [18], 17 samples from Ibrahim *et al*. [19], four samples from Rutledge *et al*. [20], one sample from Ansari *et al*. [21] and one sample from Plenderleith *et al*. [22]. Three *P. malaria-like* genomes obtained from chimpanzees (*Pan troglodytes*) were included as an outgroup [20,22]. Short read archives (SRA) from the published studies were retrieved from NCBI (accession numbers provided in S1 Table).

Short reads were trimmed to remove potential lingering adapters and preprocessed to eliminate low-quality reads (−*quality-cutoff* = 30) using the *cutadapt* program [23]. Reads shorter than 50 bp and containing “N” (*i*.*e*. ambiguous bases) were discarded (−*minimumlength* = 50 *–max-n* = 0). Cleaned paired-end reads were mapped to the *P. malariae* reference genome *PmUG01* [20] using *bwa-mem* [31] with default parameters. Duplicate reads were marked using the *MarkDuplicates* tool from the *Picard tools* v2.5.0 (broadinstitute.github.io/picard/) with default options. Local realignment around indels was performed using the *IndelRealigner* tool from *Genome Analysis Toolkit* (*GATK* [25], v3.8.0). Variants were called per sample using the *HaplotypeCaller* module in GATK with the parameter *-stand_call_conf* equals to a Phred-scaled confidence score of 10. Lastly, the different isolated variant call format (VCF) files were merged using the *GATK* module CombineGVCFs.

### Core genome SNP filtering and final dataset composition

Combining previously published dataset (n=248) with the 79 newly sequenced samples resulted in a total of 327 samples prior to filtering: 304 *P. malariae* samples, 20 *P. brasilianum* samples, and three *P. malariae-like* samples. All filtration steps are detailed in S1 Figure.

Samples with >75% missing data were removed (n=69). As several parasite strains can infect the same host, the within-host infection complexity was assessed with the *F*_*WS*_ metric [26], calculated with *vcfdo* (github.com/IDEELResearch/vcfdo; last accessed July 2022). Samples with pluri-clonal infections, *i*.*e. F*_*WS*_ ≤0.85 were removed (n=44) as in Ibrahim *et al*. [18] (S2A Fig.). For samples that are the sole representatives of their population, the *F*_*WS*_ index was replaced with a homozygosity measure to evaluate the multiplicity of infection and avoid the bias introduced by an *F*_*WS*_ value of 0 for isolated samples (S2B Fig.).

Highly related samples and clones can generate spurious signals of population structure, bias estimators of population genetic variation, and violate the assumptions of the model-based population genetic approaches used in this study [27]. The relatedness between haploid genotype pairs within each country was measured by estimating the pairwise fraction of the IBD between strains within populations using the *hmmIBD* program [28], with default parameters for recombination and genotyping error rates, and using the allele frequencies estimated by the program. Within each country, isolate pairs that shared >50% of IBD were considered highly related. In each family of related samples, only the strain with the lowest number of missing data was retained, thus removing 33 samples (S2C Fig.).

The final dataset included whole genome sequencing data of 181 samples from 29 countries, including 47 newly sequenced strains (Figure 1), and consisted in two *P. brasilianum* samples (Colombia n=1, French Guiana n=1), 179 *P. malariae* samples with four samples from South America (French Guiana n=1, Brazil n=3), eight from North Africa (Algeria n=1 and Sudan n=7), 61 from West Africa (Senegal n=1, Gambia n=1, Guinea n=5, Sierra Leone n=2, Ivory Coast n=9, Mali n=4, Ghana n=14, Togo n=1, Benin n=2, Nigeria n=24), 50 from Central Africa (Cameroon n=21, Central Africa n=4, Equatorial Guinea n=1, Gabon n=6, Congo n=11, Angola n=5), 40 from East Africa (Uganda n=14, Kenya n=10, Tanzania n=12, Malawi n=4) and 13 from Asia (India n=1, Thailand n=10, Malaysia n=1, Papua New Guinea n=1). Two *P. malariae-like* samples from Gabonese chimpanzees (*Pan troglodytes troglodytes*) were included as an outgroup. The mean sequencing depth ranged from 0.4 to 732 X for *P. malariae genomes* and from 18.4 to 521.4 X for *P. brasilianum* genomes (S1 Table).

**Figure 1.**
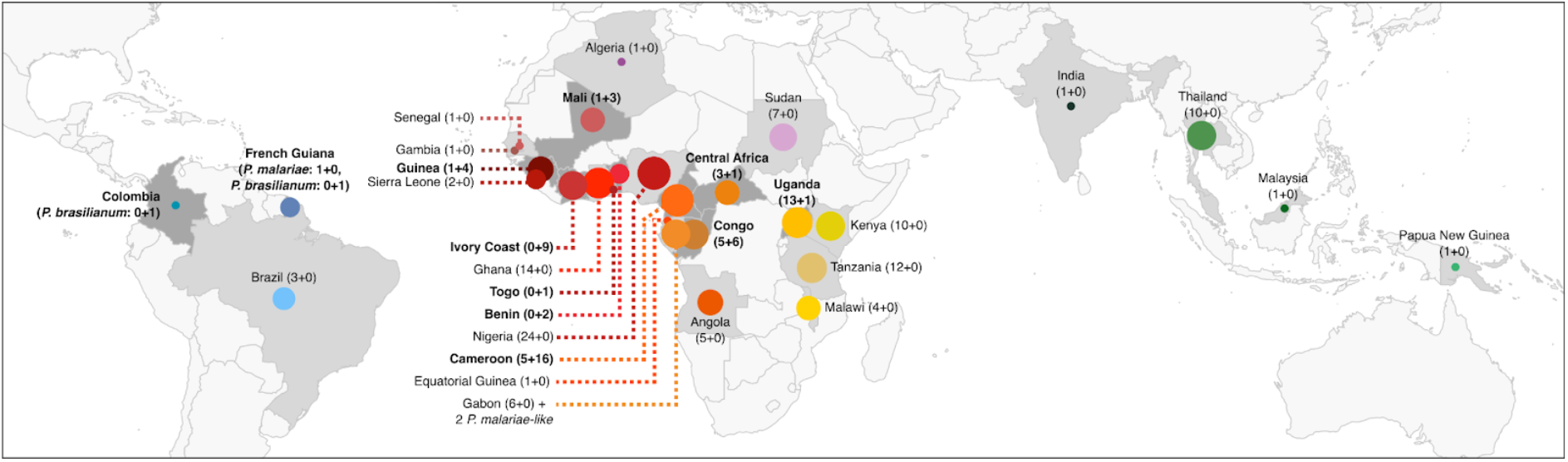
Geographic origins of the 179 *P. malariae* isolates, two *P. brasilianum*, and two *P. malariae*-like isolates from African great apes. Sample distribution by region: South America (n = 6, blue shades), North Africa (n = 8, purple shades), West Africa (n = 63, red shades), Central Africa (n = 50, orange shades), East Africa (n = 40, yellow shades), and Asia (n = 13, green shades). Sample counts are shown in brackets and with circle size. The countries involved in the sampling are shown in shades of grey. Darker shades indicate countries with newly sequenced genomes, while lighter shades indicate countries with previously published data. Bolded country names denote locations where new genomes were generated in this study. Sample counts from the literature are listed first in brackets, followed by newly sequenced samples.

Most population genetic analyses were conducted using ANGSD v0.940 [29] or ANGSD suite software, only on the core genome regions as defined by Ibrahim *et al*. [19]. All sites with a base quality ≥20 were kept. For analyses that did not require invariant sites, only SNPs with a *p-value* = 1e-6 were retained. For the maximum likelihood (ML) tree, we used a filtered VCF with only bi-allelic SNPs with ≤50% missing data, a quality score ≥30, and a minimum and maximum depth coverage set between 10X and 182X. Moreover, singletons were removed to minimize sequencing errors.

### Global population structure and genetic relationships

Principal Component Analysis (PCA) and the ancestry plots were computed with *ANGSD* and *PCAngsd* v0.98, after selecting only SNPs present in the core region of the *P. malariae* genomes and excluding SNPs with a minor allele frequency (MAF) ≤1%. The variants were linkage disequilibrium (LD) pruned to obtain a set of unlinked variants using *ngsld* v1.2 [30], with a threshold of r^2^=0.5, a window size of 5kb and a step of 1kb. After LD-pruning, 210,703 SNPs were retained for 181 individuals. The two *P. malariae-like* outgroup samples were excluded from these analyses. The ancestry plots were estimated using *PCAngsd* for *k* value (number of clusters) ranging from 2 to 16. Then, *pong* v1.5 [31] was used to analyze the PCAngsd resulting outputs and compute ancestry proportions.

The ML tree was reconstructed with IQ-TREE v2.3.4 [32] using the best-fitted model determined by ModelFinder [33]. As the dataset with outgroup comprised 118,772 SNPs of the core genome and no invariant sites, the ascertainment bias correction was added to the tested models. The best inferred model was a general time reversible (GTR) model of nucleotide evolution that integrated unequal rates and unequal base frequency. The node reliability was assessed with Ultrafast Bootstrap Approximation [34] and the SH-aLRT test [35].

### Characterization of population structure within Africa

To investigate fine-scale population structure within Africa, we restricted the dataset to African *P. malariae* isolates, as Africa represented both the most densely sampled region (n=161) and the largest proportion of newly generated genome data in this study. Because one Sudanese sample clustered with American rather than African populations, it was excluded from these subsequent analyses. The final African dataset comprised 160 samples, in 22 countries (Figure 1).

Population structure within Africa was first explored using PCA and genetic ancestry plots computed with *ANGSD* and *PCAngsd* v0.98, after selecting only SNPs present in the core region of the *P. malariae* genomes and excluding SNPs with a minor allele frequency (MAF) ≤1%. The variants were LD-pruned to obtain a set of unlinked variants using *ngsld* v1.2 [30], with a threshold of r^2^=0.5, a window size of 5kb and a step of 1kb. In total, 210,587 SNPs were used. These analyses revealed the presence of two major genetic clusters within Africa. The ancestry plots were therefore inferred with *PCAngsd* assuming K = 2.

To quantify genetic diversity within each of the two inferred African clusters, Tajima’s D [36] and nucleotide diversity (π) were measured using *pixy* v2.0.0 [37], with non-overlapping windows of 500 bp in the core genome, and only windows with ≥100 sites were kept.

Genome-wide patterns of genetic differentiation caused by reduced recombination between the two African clusters were characterized using estimates of *F*_*ST*_ and *D*_*XY*_ computed with *pixy* [37] in sliding-window of 5 kb windows and 500 bp steps. For *F*_*ST*_, only windows containing at least 100 SNPs were retained, whereas for *D*_*XY*_, windows were required to include a minimum of 1,000 sites. Genomic regions proximal to telomeres and centromeres exhibited elevated values due to edge effects (incomplete windows). Since the core genome used has only been defined on a few genomes [19], it is possible that this pattern is also due to the inclusion of telomeric or subtelomeric regions in the windows. The windows were therefore excluded from subsequent analyses. However, the patterns found in the genome with these two metrics might be caused by a reduction in recombination rate between the two clusters or because of changes in population demographic history, and may not necessarily result from positive selection [38]. Thus, lineage-specific tests of selection were subsequently performed.

### Lineage specific positive selection detection

To identify genomic regions potentially under positive selection in each African cluster, the population branch-site (PBS) values [39] were calculated for each cluster, with *pixy* [37] using sliding windows of 5 kb and a step size of 500 bp. The outgroup populations were the alternative African cluster and Thai population. All windows with <100 SNPs were removed to avoid extreme values caused by a low SNP number. The 0.1% most extreme values were considered evidence that the genomic regions displayed signs of selection in the African cluster. Once positive selection signals were detected, the identified genes were annotated using the general feature format (GFF) file *GCF_900090045*.*1_PmUG01_genomic*.*gff* (available from PlasmoDB) and the *intersect* function of *BEDtools* v2.31.1 [40]. Additional information was retrieved from *PlasmoDB* (plasmodb.org, accessed in December 2025).

## Results

### *Plasmodium brasilianum* infections in South American NHPs

Among 226 samples collected from NHPs across two Latin American countries (14 Colombian, and 212 French Guianese), 20 (n=4/14 from Colombia, and n=16/212 from French Guiana), were positive for *Plasmodium* using the *cytochrome-b* based PCR assay [16] (Table S1). Twenty *Plasmodium*-infected samples were identified across six NHP species (*Alouatta macconnelli* (n = 10/140), *Saguinus midas* (n = 3/58), *Ateles hybridus* (n = 1/5), *Alouatta seniculus* (n = 2/5), *Aotus griseimembra* (n = 1), and *Cebus versicolor* (n = 1/3)). Sequence analyses showed that all infections were attributable to *P. brasilianum*. All positive samples were then subjected to selective whole-genome amplification (sWGA) followed by whole-genome sequencing. However, after applying stringent quality filtering and SNP selection criteria, only two *P. brasilianum* samples (one from a Colombian *A. seniculus* and one from a French Guianan *A. hybridus*) retained sufficient high-quality SNPs to be included in the final dataset.

### Cryptic population structure within African populations

To explore the genetic relationships between *P. malariae* populations and with *P. brasilianum*, the population structure was characterized using complementary population genomic approaches, including principal component analysis (PCA), maximum likelihood (ML) phylogenetic tree, and model-based individual genetic ancestry inference.

The first three principal components (PCs) of the PCA revealed four major genetic clusters (Fig. 2A). PC1 separated African isolates into two distinct clusters, whereas PC2 distinguished Asian populations from all others. Along PC3, South American *P. malariae* and *P. brasilianum* samples were separated from the remaining populations. Although the Asian and American clusters were clearly distinguishable in the ML tree (Fig. 2B), the two African clusters were not. Overall, few nodes showed sufficient statistical support to be considered reliable (using thresholds established in the literature [34,35,41]), with the exception of those within the Asian and American clades. Furthermore, the two isolates of *P. brasilianum* were connected basally to the Latin American clade, although two samples of Brazilian *P. malariae* (*PM_BRA_004* and *PM_BRA_007*) were actually included in the African diversity and not the Latin American (respectively ✴ and ✳ on Fig. 2B). One of the isolates (*PM_BRA_004*) is even very similar to a Nigerian isolate (*PM_NGA_029*), which can be seen on the ML Tree and PCA (✴ on Fig. 2, and S3 Fig.). Moreover, one Sudanese sample (*PM_SDN_009*) is genetically close to the American *P. malariae* clade (፠ on Fig. 2).

**Figure 2.**
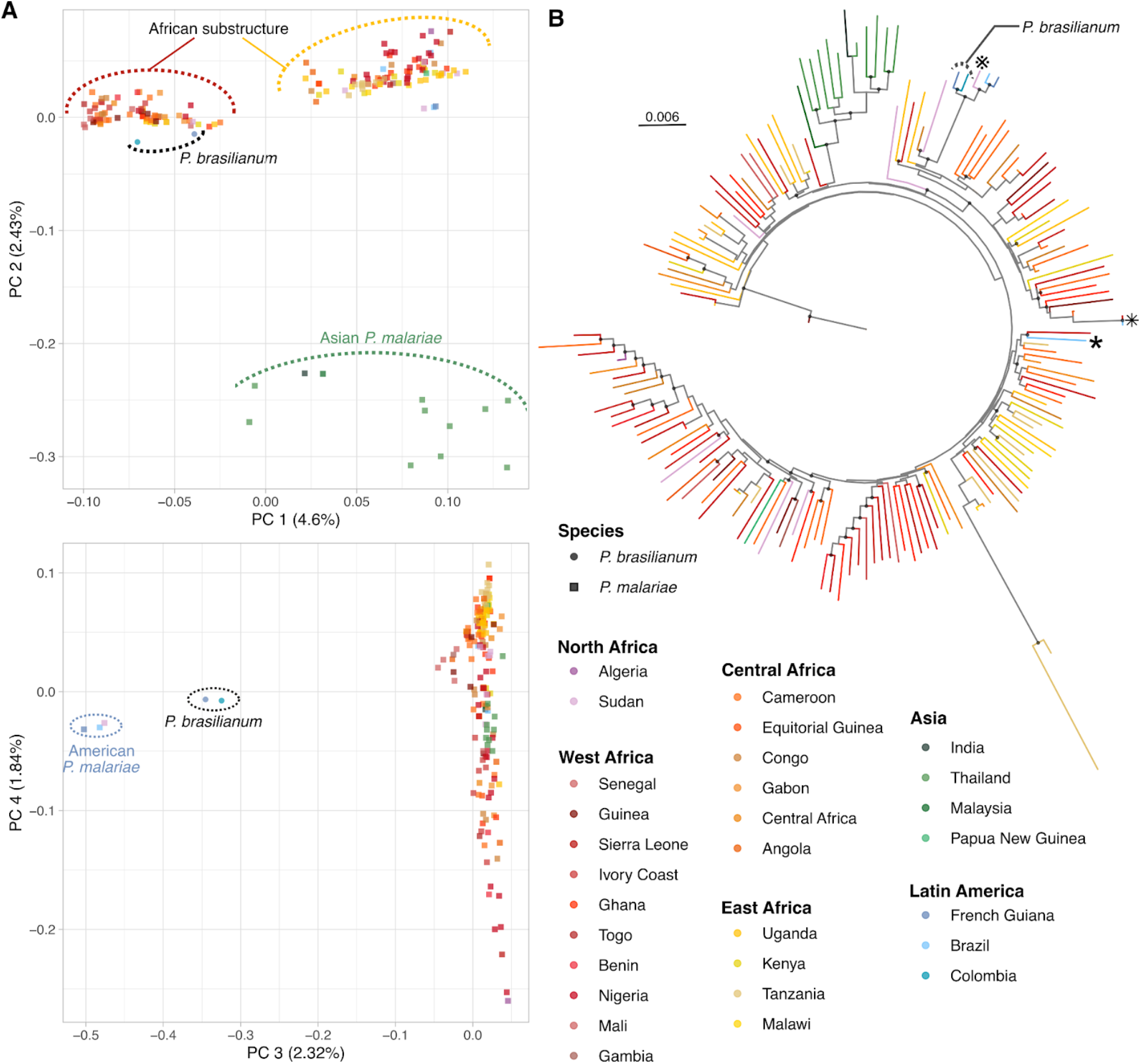
*P. malariae* and *P. brasilianum* genetic structure. (A) Principal component (PC) analysis of 179 *P. malariae* and two *P. brasilianum* strains showing the first, second and third PCs based on the genotype likelihood of 210,703 unlinked SNPs. *Plasmodium brasilianum* are indicated by circles and *P. malariae* by squares. (B) Maximum likelihood phylogenetic tree of 179 *P. malariae*, two *P. brasilianum* and two *P. malariae-like* individuals. The tree includes two *P. malariae-like* strains used as outgroups. Note that the length of the outgroup branch (central line) has been truncated. Black dots at some nodes indicate highly supported nodes (both SH-aLRT ≥ 80% and UFboot ≥ 95%, following the thresholds established in the literature [34,35,41]). The symbols (፠,✴, and ✳) mark samples whose placements are discussed in the paper.

The genetic ancestry analyses were consistent with the PCA and the ML phylogeny. As observed in the PCA and the ML tree, the Asian and American clusters were well defined, emerging from *K* = 3 for the Asian cluster and *K* = 5 for the American cluster (Fig. 3 and S4). In addition, two Brazilian *P. malariae* samples clearly clustered with African populations (✴ and ✳ on Fig. 3A), in agreement with the PCA and ML tree. Similarly, the Sudanese sample appeared to be genetically closer to the American cluster (፠ on Fig. 3A).

**Figure 3.**
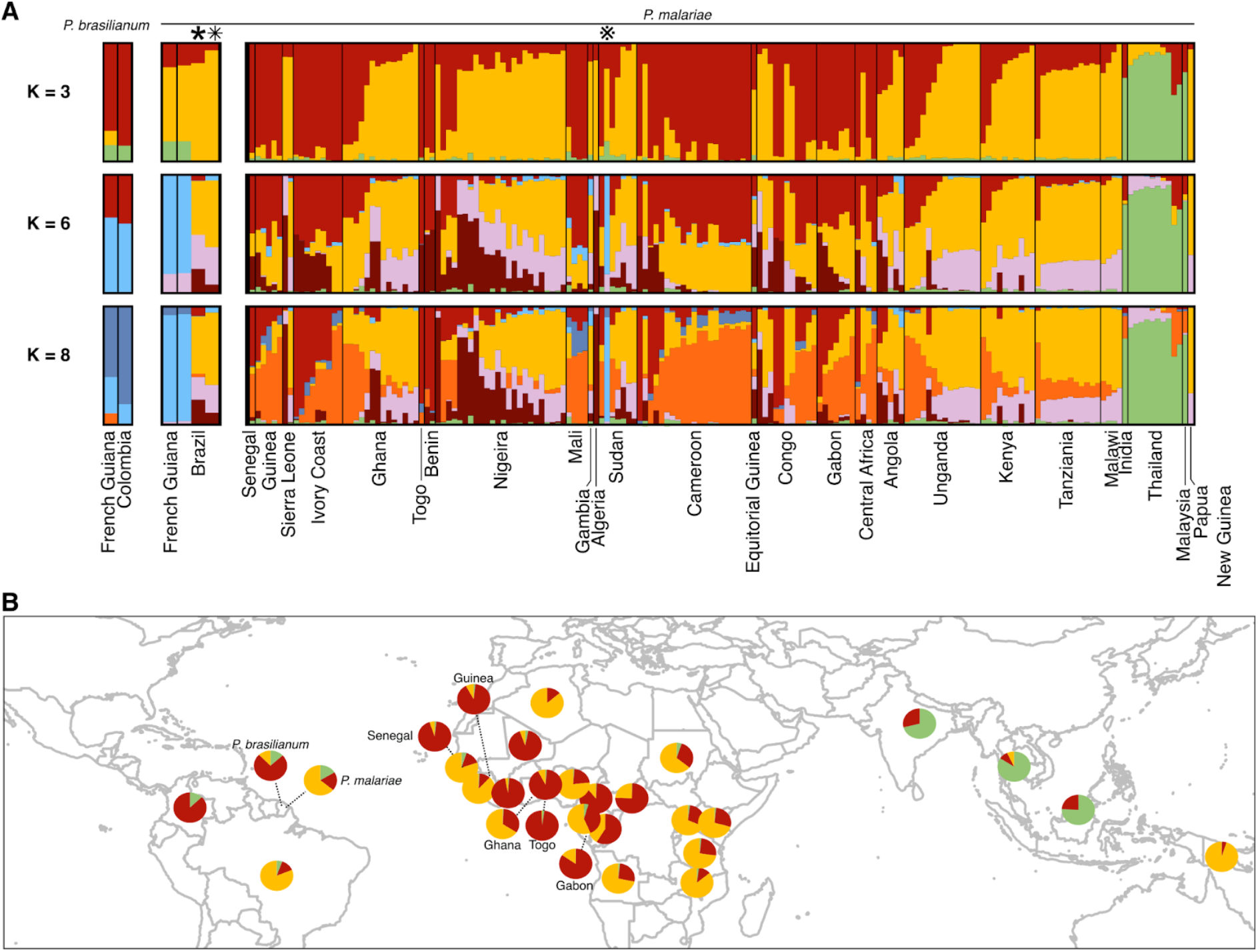
Ancestry of *P. malariae* and *P. brasilianum*. (A) Individual genetic ancestry assuming K=3, K=6 and K=8 genetic clusters estimated using *PCAngsd* (see Fig. S4 for the other K values). K=6 is the best K according to the method of Meisner and Albrechtsen [42] (see Fig. S3). The symbols (፠,✳, and ✴) mark samples whose placements are discussed in the paper. (B) Individual genetic ancestry assuming K=3 genetic clusters estimated using *PCAngsd* displayed as pie charts on the world map.

At a finer scale within Africa, two clusters were also detected (Fig. 3). However, they were less clearly defined than in the PCA (Fig. 2A), as many isolates showed admixed genetic ancestry. This admixture may explain the lack of clear separation between the African clusters in the ML tree, as well as the low statistical support observed for the nodes within the African diversity (Fig. 2B). Moreover, no clear geographic structuring was apparent between the two African clusters (Fig. 2B), which appear to coexist in the same regions and recombine. Because no obvious biological or geographic criteria allow these clusters to be distinguished, they were referred hereafter according to the colors used in the ancestry plots: cluster R (red) and cluster Y (yellow).

Finally, *P. brasilianum* isolates seem genetically closer to cluster R, whereas American *P. malariae* samples seemed more closely associated with cluster Y (Fig. 3).

### Genomic evidence for divergence and selection in the two sympatric cryptic African clusters

Although all isolates originated from the same continent, two genetic clusters were identified within African *P. malariae* populations (Fig. 2 and 3). We therefore investigated the presence of genetic substructure within African *P. malariae*. Because one Sudanese sample (*PM_SDN_009*) clustered with American rather than African populations (Figs. 2 and 3), it was excluded from subsequent analyses. A PCA restricted to African samples (n=160) confirmed the separation into two distinct genetic clusters (Fig. 4A). In contrast, ancestry inference using *PCAngsd* revealed a less sharply defined pattern, with several individuals exhibiting mixed ancestry between the two clusters. Despite the absence of a clear geographic partitioning, cluster Y was more prevalent in East and North Africa, whereas cluster R was more frequently observed in West Africa (Fig. 4B). Notably, isolates from both clusters were detected across multiple regions of Africa, indicating broad geographic overlap.

**Figure 4.**
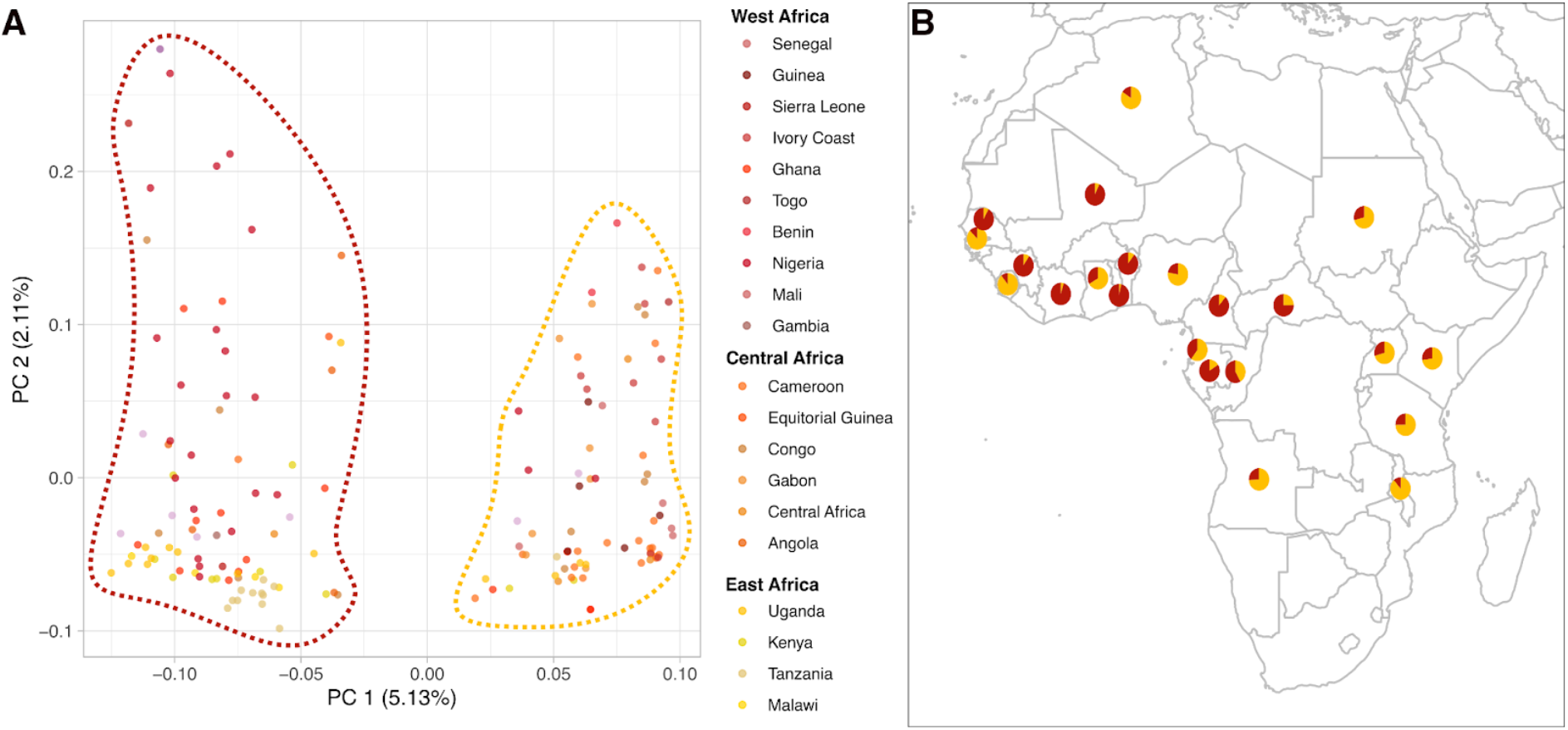
Genetic characterization of two *P. malariae* clusters in Africa. (A) Principal component analysis (PCA) of 160 African *P. malariae* isolates, showing the first two principal components derived from genotype likelihoods at 210,587 unlinked SNPs. (B) Individual genetic ancestry of 160 African *P. malariae* strains assuming K=2 genetic clusters estimated using *PCAngsd*, represented as pie charts on a map of Africa.

To further characterize the two African clusters, we retained only samples predominantly associated with a single cluster, defined as individuals showing more than 70% ancestry for one cluster in the ancestry analysis. This resulted in 74 samples in cluster R and 70 samples in cluster Y. The genome-wide distributions of nucleotide diversity (π) and Tajima’s D [36] differed significantly between the two clusters (Wilcoxon signed-rank tests, *p-value* < 0.001 for both statistics). However, a Wilcoxon effect size test [43] revealed that the difference was very small, even negligible (*r* < 0.1) [44]. Specifically, π was slightly higher (medians of 2.16×10^-4^ for cluster Y and 3.09×10^-4^ for cluster R) and Tajima’s D was marginally lower (medians of -1.107 for cluster Y and -1.163 for cluster R) in cluster R than in cluster Y (Fig. S5).

To better understand the origin of these two clusters and identify genomic features underlying their differentiation, we performed genome-wide scans of *F*_*ST*_ and *D*_*XY*_. Genomic regions where the recombination is less efficient between lineages, named islands of differentiation, are expected to exhibit elevated values of both *F*_ST_ and D_XY_ [45,46]. No region showed high values for both metrics. However, numerous *F*_ST_ peaks suggest differentiation driven by selection specific to each lineage (Fig. S6).

Given the genetic differentiation between the two African clusters, they may be subject to distinct selective pressures. We therefore assessed cluster-specific signals of adaptation by estimating the population branch statistic (*PBS*), an *F*_*ST*_-based metric, for each African cluster using comparisons with the alternate African cluster and a Thai population. Windows with outlier *PBS* values (top 0.1%) were interpreted as regions showing elevated differentiation specific to the focal African cluster. We identified 489 outlier windows in the cluster Y and cluster R, corresponding to positive selection in the CDS of 203 genes (cluster Y) and 112 genes (cluster R) (Figure 5 and S2 Table).

**Figure 5.**
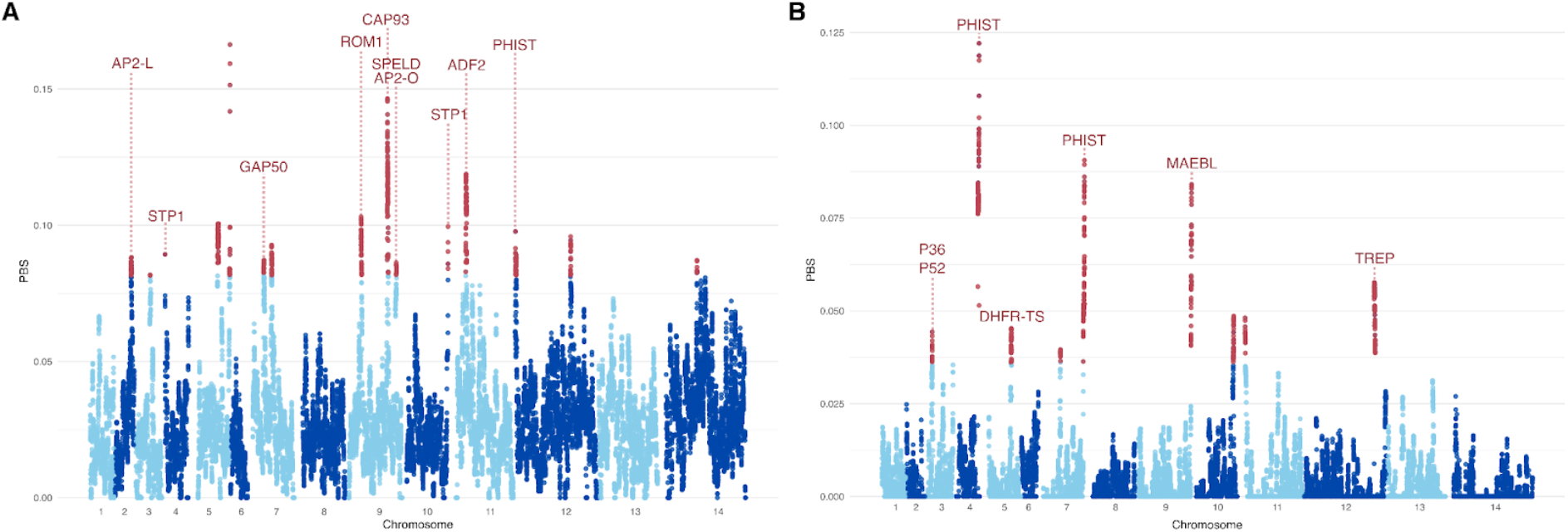
Evidence of selection signals in African *P. malariae*. **(A & B)** Manhattan plots of population branch statistic (PBS) values, an *F*_*ST*_-like statistic that measures allele frequency differentiation relative to two reference populations. For each African *P. malariae* cluster, we compared it with the other African cluster and Thai samples as outgroups. Red points indicate the top 0.1% of PBS values, suggesting higher allelic frequency differentiation compared to the reference populations, and thus indicating significant evidence of positive selection that affects these regions. A corresponds to Cluster Y and B to Cluster R.

More specifically, 22 of the identified genes are known or predicted to be involved in interactions between the parasite and its hosts (in bold in S2 Table). Genes putatively associated with interactions with primate hosts, including *AP2-L* [47], *ROM1* [48], *SPELD* [49], and *STP1* [50] showed signals of selection in cluster Y, whereas 6-cysteine proteins (*P36* and *P52* [51–53]) appeared to be under selection in cluster R. In addition, several genes displaying signatures of selection are implicated in interactions with the mosquito vector, including *AP2-O* [54] and *CAP93* [55] in cluster Y and *TREP* [56] in cluster R. Two genes showing signatures of selection in cluster Y, *GAP50* and *ADF2*, are predicted to be involved in interactions with both primate and mosquito hosts [57,58]. Only one annotated gene implicated in interactions in both hosts, *MAEBL* [59,60], appeared to be under selection in cluster R. Notably, members of the *PHIST* gene family exhibited signals of selection in both clusters. This family has been implicated in a range of biological processes, including cytoadherence, gametocytogenesis, host cell modification, and the production of extracellular vesicles [61,62]. However, none of the *PHIST* genes identified here have been functionally characterized in *P. malariae* (S2 Table). Furthermore, signals of selection at the *DHFR-TS* gene, which is associated with resistance to antimalarial treatment in *Plasmodium* [63,64], were detected exclusively in cluster R (Fig. 5B). In total, we identified 315 genes exhibiting signatures of positive selection in African *P. malariae*, including multiple candidates involved in interactions with both vertebrate hosts and mosquito vectors.

## Discussion

Although *P. malariae* is a relatively overlooked malaria parasite, recent efforts have substantially expanded the availability of whole-genome data [18–22]. The present study makes a significant contribution to the ongoing corpus by sequencing 59 samples of *P. malariae* and 20 samples of *P. brasilianum* from NHPs. The *Plasmodium* screening performed on 297 NHP specimens revealed active circulation of *P. brasilianum* across multiple primate species in Colombia and French Guiana, highlighting the persistence of this parasite in sylvatic transmission cycles. Following stringent data curation, 179 high-quality *P. malariae* (including 160 from Africa) and only two *P. brasilianum* genomes were retained. This substantial representation of African *P. malariae* isolates allowed us to conduct a global analysis of parasite diversity while providing detailed insights into population structure and evolutionary dynamics within Africa, where the species is the most prevalent.

### Circulation of *P. brasilianum* in NHPs

A total of 226 American NHPs from Colombia (n = 14) and French Guiana (n = 212) were screened for *Plasmodium* infection. *Cytochrome-b* PCR screening identified 20 *P. brasilianum* infected individuals (8.9%), including four from Colombia and sixteen from French Guiana. Infections were detected across six host species: *A. macconnelli* (10/140), *S. midas* (3/58), *A. hybridus* (1/5), *A. seniculus* (2/5), *A. griseimembra* (1/1), and *C. versicolor* (1/3). No infections were detected in the remaining four screened species (*A. paniscus, S. apella, S. sciureus and P. pithecia)* but this may be due to the small sample size for these species (from n=1 to n=7). The detection of infections across multiple primate genera, including spider and howler monkeys, as well as smaller-bodied taxa, is consistent with previous reports indicating that *P. brasilianum* circulates among phylogenetically diverse American primates [7–9,11]. Although sample sizes varied among species, the presence of infection in several host lineages further supports the existence of a multi-host sylvatic transmission cycle within forest ecosystems.

Despite the successful detection of infections, the recovery of adequate genomic data has proven challenging. After sWGA and stringent SNP filtering, only two *P. brasilianum* genomes were retained for population genomic analyses (S1 Fig. and Table S1). This limitation likely reflects the typically low parasitemia observed in natural infections and high host DNA contamination, underscoring the technical challenges inherent to studying malaria parasites in wildlife hosts. Nevertheless, the positioning of these two *P. brasilianum* genomes within the broader *P. malariae* diversity confirms their close genetic relationship with American *P. malariae* isolates. Moreover, the affinity of *P. brasilianum* with African cluster R suggests possible historical connectivity between African and American parasite populations, potentially linked to past human-mediated dispersal events. Nevertheless, the limited number of high-quality *P. brasilianum* genomes currently available constrains inference. Future studies should prioritize the generation of well-covered genomes from diverse American monkeys and geographic regions to clarify the origin, diversification, and timing of introduction of this lineage in Latin America.

### Hidden structure in Africa: discovery of two recombinant cryptotypes of *P. malariae*

At a global scale, we identified two well-defined genetic groups corresponding to Asia and South America (Fig. 2A and B), consistent with the literature [18]. In contrast, our study revealed a previously unnoticed population structure within Africa. While Ibrahim *et al*. [18] noted a potential substructure among African isolates, this pattern was not formally supported. The discrepancy between these two studies likely reflects methodological differences. In particular, the removal of closely related samples in our dataset reduced the risk of inflated structure signals and biased estimators of genetic variation, which can arise from uneven sampling of related infections [27], a step that was incorporated in the present study.

To our knowledge, this study provides the first evidence for the presence of two distinct genetic clusters within African *P. malariae*, designated as clusters R and Y (Fig. 2, 3 and 4). These two clusters appear to coexist in sympatry and recombine across multiple African regions, with no obvious geographic structuring (Fig. 4B). This pattern is reminiscent of the “cryptic” population structure recently described in *P. falciparum*, where specific genetic backgrounds can be detected across Africa despite extensive recombination and broad geographic overlap. For instance, Miotto *et al*. [15] identified a low-frequency, continent-wide *P. falciparum* cluster (*AF1*) characterzed by a shared multi-locus genetic background, termed a cryptotype, that persists despite outbreeding with local parasites. More broadly, African *P. falciparum* also shows macro-scale population structure (West/Central/East) revealed by genome-wide clustering [65,66]. One possible (but currently untested) explanation for their distribution could be ecological heterogeneity, such as the presence of tropical forest-associated transmission systems [67,68]. Indeed, cluster R seems more prevalent in countries covered by tropical forests (*e*.*g*: Gabon, Cameroon, Central Africa, Benin, Togo, Ivory Coast and Guinea). However, testing this hypothesis would require more finely resolved sampling, as our dataset primarily consists of isolates obtained from travelers visiting these African countries, for which geographic information about infection is limited to country-level. Furthermore, our sampling covers several years (S1 Table), and the tropical forest coverage highly transformed into tropical savanna changed between 2010 and 2017 [69], which further complicates direct ecological interpretations. A better understanding of how parasites shift their distribution and move across populations could greatly contribute also from a One Health perspective.

This hypothesis is consistent with our observation that *P. brasilianum* samples are genetically closer to cluster R, whereas American *P. malariae* samples are more closely related to cluster Y (Fig. 2A and 3). Because *P. brasilianum* is adapted to NHPs, its transmission might be sylvatic [12]. This pattern might therefore reflect adaptation to forest-associated environments, maybe with a different transmission dynamic, although further ecological and entomological data would be required to evaluate this hypothesis.

### African cluster-specific adaptation signals are dominated by host-interaction genes

Genome-wide scans aimed at identifying regions showing cluster-specific signals of positive selection did not reveal genes with clear links to environmental adaptation (Fig. 4C). However, our results point toward processes more closely tied to within-host biology and host–parasite interactions rather than adaptation to external abiotic conditions.

We identified multiple genes showing signatures of selection that are putatively involved in interactions with hosts, including both mosquito vectors and vertebrate hosts, with patterns that differ between the African clusters (Fig. 5 and S2 Table). Although many candidate loci remain functionally unknown in *P. malariae*, their putative roles have been extrapolated by orthology-based inferences from studies in other *Plasmodium* species infecting primates (*P. falciparum, P. knowlesi*) or rodents (*P. berghei*). Among genes without functional annotation in *Plasmodium*, some genes may have played a role in African *P. malariae* adaptation to the mosquito and/or primate host. Thus, functional follow-up, as experimental validation will be essential to move from statistical signatures to mechanistic understanding of *P. malariae* adaptation

In cluster R, the *DHFR-TS* gene shows signs of selection. This signal has been previously reported and functionally validated in *P. malariae* [18]. Indeed, as in *P. falciparum*, this gene is involved in resistance to antimalarial treatment [18,64,70]. Importantly, this represents the only selection signal associated with drug resistance detected in our study which is also the sole gene under selection in Africa that overlaps with findings reported by Ibrahim *et al*. [18]. Differences between the two studies may reflect both analytical and conceptual choices. Ibrahim *et al*. [18] employed approaches based on identity-by-descent and haplotype length, whereas our analyses relied on the PBS, which is derived from allele frequency differentiation (*F*_*ST*_). Haplotype-based methods are generally more sensitive to recent selection but may fail to detect older selective events due to the breakdown of linkage disequilibrium through recombination. Our approach may capture older selection signals. In addition, Ibrahim *et al*. [18] treated African *P. malariae* as a single population, whereas our analyses explicitly accounted for the presence of two distinct African genetic clusters and focused on cluster-specific signals of adaptation.

In conclusion, although *P. malariae* remains a neglected malaria parasite, it poses a tangible public health risk and represents an additional challenge to malaria control and elimination efforts. By assembling and analyzing the largest and most comprehensive whole-genome dataset for this species to date (to our knowledge), we provide new insights into the global genetic diversity of *P. malariae*. The identification of two recombinant and sympatric African genetic clusters challenges current views of *P. malariae* transmission and diversity across the continent. This complexity is particularly relevant given that each cluster exhibits distinct signatures of adaptation to human hosts and mosquito vectors, even though the ecological or evolutionary drivers of their divergence still remain unclear. Our findings underscore the need for larger, geographically finer-scale sampling to better resolve the evolutionary dynamics of *P. malariae*. Indeed, hidden population substructure can hinder malaria control strategies by obscuring transmission pathways, masking local adaptation, and potentially influencing responses to interventions such as drug treatment or vector control. Thus, by revealing previously unrecognized population structure in Africa, this study highlights the importance of integrating population genomic data into malaria surveillance, control, and elimination programs to ensure that control and elimination strategies account for the full diversity of malaria parasites circulating in endemic regions.

## Acknowledgments

The bioinformatics analyses were performed on the Core Cluster of the Institut Français de Bioinformatique (IFB) (ANR-11-INBS-0013). The authors would like to thank the members of the network for their participation in the surveillance of imported malaria in France, and the staff of the National Reference Center for Malaria, who made it possible to establish the biological collection.

## Funding

JCJC GENAD, ANR-20-CE35-0003 (VR)

MICETRAL, ANR-19-CE350010 (FP)

project ORA in the LabEx CEBA, ANR-10-LABX-25-01 (VR)

the United States (U.S.) Agency for International Development and the U.S. Centers for Disease Control and Prevention (CDC) for the project entitled “Evaluation of ZIKV potential to establish a sylvatic transmission cycle in Colombia” (CG, SR, AL)

## Author contributions

Conception: V.R., F.P., M.C.F. and M.J.M.L.

Funding acquisition: V.R., F.P., M.C.F.

Biological data acquisition and management: V.R., S.H., B.d.T., C.G., S.R, A.L.

Sequence data acquisition: C.A.

Method development and data analysis: M.J.M.L., A.P., V.R., F.P., and M.C.F.

Interpretation of the results: V.R., F.P., M.C.F., and M.J.M.L.

Drafting of the manuscript: M.J.M.L., V.R., M.C.F., and F.P.

Reviewing and editing of the manuscript: V.R., F.P., M.C.F., and M.J.M.L. with the contribution from all the co-authors

## Data availability

The sequences of the newly sequenced *P. malariae* and *P. brasilianum* samples have been deposited on NCBI (Bioproject PRJNA1168452). SNP data files (VCF format) produced in this study as well as the related scripts are available in DataSuds repository (IRD, France) at https://doi.org/10.23708/PXTDVF.

The scripts used in this study are also available in this github repository: https://github.com/MargauxLefebvre/WGS_Pmalariae.git

## References

1. Geneva: World Health Organization. World malaria report 2025: addressing the threat of antimalarial drug resistance. 2025.

2. Rougeron V, Boundenga L, Arnathau C, Durand P, Renaud F, Prugnolle F. A population genetic perspective on the origin, spread and adaptation of the human malaria agents Plasmodium falciparum and Plasmodium vivax. FEMS Microbiol Rev. 2021. doi:10.1093/femsre/fuab047

3. Hawadak J, Dongang Nana RR, Singh V. Global trend of Plasmodium malariae and Plasmodium ovale spp. malaria infections in the last two decades (2000–2020): a systematic review and meta-analysis. Parasit Vectors. 2021;14: 297. doi:10.1186/s13071-021-04797-0

4. Sutherland CJ. Persistent Parasitism: The Adaptive Biology of Malariae and Ovale Malaria. Trends Parasitol. 2016;32: 808–819. doi:10.1016/j.pt.2016.07.001

5. Collins WE, Jeffery GM. Plasmodium malariae: parasite and disease. Clin Microbiol Rev. 2007;20: 579–592.

6. Langford S, Douglas NM, Lampah DA, Simpson JA, Kenangalem E, Sugiarto P, et al. Plasmodium malariae Infection Associated with a High Burden of Anemia: A Hospital-Based Surveillance Study. PLoS Negl Trop Dis. 2016;9: e0004195. doi:10.1371/journal.pntd.0004195

7. Alvarenga DAM, Pina-Costa A, Bianco C, Moreira SB, Brasil P, Pissinatti A, et al. New potential Plasmodium brasilianum hosts: tamarin and marmoset monkeys (family Callitrichidae). Malar J. 2017;16: 71. doi:10.1186/s12936-017-1724-0

8. Fandeur T, Volney B, Peneau C, De Thoisy B. Monkeys of the rainforest in French Guiana are natural reservoirs for P. brasilianum/P. malariae malaria. Parasitology. 2000/01/01 ed. 2000;120: 11–21. doi:10.1017/S0031182099005168

9. Rondón S, León C, Link A, González C. Prevalence of Plasmodium parasites in non-human primates and mosquitoes in areas with different degrees of fragmentation in Colombia. Malar J. 2019;18: 276. doi:10.1186/s12936-019-2910-z

10. Voinson M, Nunn CL, Goldberg A. Primate malarias as a model for cross-species parasite transmission. Wesolowski A, Perry GH, editors. eLife. 2022;11: e69628. doi:10.7554/eLife.69628

11. Fuentes-Ramírez A, Jiménez-Soto M, Castro R, Romero-Zuñiga JJ, Dolz G. Molecular Detection of Plasmodium malariae/Plasmodium brasilianum in Non-Human Primates in Captivity in Costa Rica. PLOS ONE. 2017;12: e0170704. doi:10.1371/journal.pone.0170704

12. Lalremruata A, Magris M, Vivas-Martínez S, Koehler M, Esen M, Kempaiah P, et al. Natural infection of Plasmodium brasilianum in humans: man and monkey share quartan malaria parasites in the Venezuelan Amazon. EBioMedicine. 2015;2: 1186–1192.

13. Yman V, Wandell G, Mutemi DD, Miglar A, Asghar M, Hammar U, et al. Persistent transmission of Plasmodium malariae and Plasmodium ovale species in an area of declining Plasmodium falciparum transmission in eastern Tanzania. PLoS Negl Trop Dis. 2019;13: e0007414. doi:10.1371/journal.pntd.0007414

14. Bajic M, Ravishankar S, Sheth M, Rowe LA, Pacheco MA, Patel DS, et al. The first complete genome of the simian malaria parasite Plasmodium brasilianum. Sci Rep. 2022;12: 19802. doi:10.1038/s41598-022-20706-6

15. Miotto O, Amambua-Ngwa A, Amenga-Etego LN, Abdel Hamid MM, Adam I, Aninagyei E, et al. Identification of complex Plasmodium falciparum genetic backgrounds circulating in Africa: a multicountry genomic epidemiology analysis. Lancet Microbe. 2024;5. doi:10.1016/j.lanmic.2024.07.004

16. Prugnolle F, Durand P, Neel C, Ollomo B, Ayala FJ, Arnathau C, et al. African great apes are natural hosts of multiple related malaria species, including Plasmodium falciparum Proc Natl Acad Sci. 2010;107: 1458–1463.

17. Ben-Rached F, Subudhi AK, Li C, Alawi M, Satyam R, Xu S, et al. Leveraging genomic insights from the neglected malaria parasites P. malariae and P. ovale using selective whole genome amplification (SWGA) approach. BMC Genomics. 2025;26: 118. doi:10.1186/s12864-025-11292-8

18. Ibrahim A, Mohring F, Manko E, van Schalkwyk DA, Phelan JE, Nolder D, et al. Genome sequencing of Plasmodium malariae identifies continental segregation and mutations associated with reduced pyrimethamine susceptibility. Nat Commun. 2024;15: 10779. doi:10.1038/s41467-024-55102-3

19. Ibrahim A, Diez Benavente E, Nolder D, Proux S, Higgins M, Muwanguzi J, et al. Selective whole genome amplification of Plasmodium malariae DNA from clinical samples reveals insights into population structure. Sci Rep. 2020;10: 10832. doi:10.1038/s41598-020-67568-4

20. Rutledge GG, Böhme U, Sanders M, Reid AJ, Cotton JA, Maiga-Ascofare O, et al. Plasmodium malariae and P. ovale genomes provide insights into malaria parasite evolution. Nature. 2017;542: 101–104. doi:10.1038/nature21038

21. Ansari HR, Templeton TJ, Subudhi AK, Ramaprasad A, Tang J, Lu F, et al. Genome-scale comparison of expanded gene families in Plasmodium ovale wallikeri and Plasmodium ovale curtisi with Plasmodium malariae and with other Plasmodium species. Int J Parasitol. 2016;46: 685–696. doi:10.1016/j.ijpara.2016.05.009

22. Plenderleith LJ, Liu W, Li Y, Loy DE, Mollison E, Connell J, et al. Zoonotic origin of the human malaria parasite Plasmodium malariae from African apes. Nat Commun. 2022;13: 1868. doi:10.1038/s41467-022-29306-4

23. Martin M. Cutadapt removes adapter sequences from high-throughput sequencing reads. EMBnet J. 2011;17: 10–12.

24. Li H, Durbin R. Fast and accurate short read alignment with Burrows–Wheeler transform. bioinformatics. 2009;25: 1754–1760.

25. McKenna A, Hanna M, Banks E, Sivachenko A, Cibulskis K, Kernytsky A, et al. The Genome Analysis Toolkit: a MapReduce framework for analyzing next-generation DNA sequencing data. Genome Res. 2010;20: 1297–1303.

26. Amegashie EA, Amenga-Etego L, Adobor C, Ogoti P, Mbogo K, Amambua-Ngwa A, et al. Population genetic analysis of the Plasmodium falciparum circumsporozoite protein in two distinct ecological regions in Ghana. Malar J. 2020;19: 437. doi:10.1186/s12936-020-03510-3

27. Wang J. Effects of sampling close relatives on some elementary population genetics analyses. Mol Ecol Resour. 2018;18: 41–54. doi:10.1111/1755-0998.12708

28. Schaffner SF, Taylor AR, Wong W, Wirth DF, Neafsey DE. hmmIBD: software to infer pairwise identity by descent between haploid genotypes. Malar J. 2018;17: 196. doi:10.1186/s12936-018-2349-7

29. Korneliussen TS, Albrechtsen A, Nielsen R. ANGSD: Analysis of Next Generation Sequencing Data. BMC Bioinformatics. 2014;15: 356. doi:10.1186/s12859-014-0356-4

30. Fox EA, Wright AE, Fumagalli M, Vieira FG. ngsLD: evaluating linkage disequilibrium using genotype likelihoods. Bioinformatics. 2019;35: 3855–3856. doi:10.1093/bioinformatics/btz200

31. Behr AA, Liu KZ, Liu-Fang G, Nakka P, Ramachandran S. pong: fast analysis and visualization of latent clusters in population genetic data. Bioinformatics. 2016;32: 2817–2823. doi:10.1093/bioinformatics/btw327

32. Nguyen LT, Schmidt HA, von Haeseler A, Minh BQ. IQ-TREE: A fast and effective stochastic algorithm for estimating maximum likelihood phylogenies. Mol Biol Evol. 2015;32: 268–274. doi:10.1093/molbev/msu300

33. Kalyaanamoorthy S, Minh BQ, Wong TKF, von Haeseler A, Jermiin LS. ModelFinder: fast model selection for accurate phylogenetic estimates. Nat Methods. 2017;14: 587–589. doi:10.1038/nmeth.4285

34. Hoang DT, Chernomor O, von Haeseler A, Minh BQ, Vinh LS. UFBoot2: Improving the Ultrafast Bootstrap Approximation. Mol Biol Evol. 2018;35: 518–522. doi:10.1093/molbev/msx281

35. Guindon S, Dufayard J-F, Lefort V, Anisimova M, Hordijk W, Gascuel O. New Algorithms and Methods to Estimate Maximum-Likelihood Phylogenies: Assessing the Performance of PhyML 3.0. Syst Biol. 2010;59: 307–321. doi:10.1093/sysbio/syq010

36. Tajima F. Statistical method for testing the neutral mutation hypothesis by DNA polymorphism. Genetics. 1989;123: 585.

37. Korunes KL, Samuk K. pixy: Unbiased estimation of nucleotide diversity and divergence in the presence of missing data. Mol Ecol Resour. 2021;21: 1359–1368. doi:10.1111/1755-0998.13326

38. Cruickshank TE, Hahn MW. Reanalysis suggests that genomic islands of speciation are due to reduced diversity, not reduced gene flow. Mol Ecol. 2014;23: 3133–3157. doi:10.1111/mec.12796

39. Yi X, Liang Y, Huerta-Sanchez E, Jin X, Cuo ZXP, Pool JE, et al. Sequencing of 50 human exomes reveals adaptation to high altitude. science. 2010;329: 75–78.

40. Quinlan AR, Hall IM. BEDTools: a flexible suite of utilities for comparing genomic features. Bioinformatics. 2010;26: 841–842. doi:10.1093/bioinformatics/btq033

41. Minh BQ, Nguyen MAT, von Haeseler A. Ultrafast Approximation for Phylogenetic Bootstrap. Mol Biol Evol. 2013;30: 1188–1195. doi:10.1093/molbev/mst024

42. Meisner J, Albrechtsen A. Inferring Population Structure and Admixture Proportions in Low-Depth NGS Data. Genetics. 2018;210: 719–731. doi:10.1534/genetics.118.301336

43. Rosenthal R, Cooper H, Hedges L. Parametric measures of effect size. Handb Res Synth. 1994;621: 231–244.

44. Fiel Peres F. Effect sizes for nonparametric tests. Biochem Medica. 2026;36: 5–16.

45. Liu Y, Yu W, Wu B, Li J. Patterns of genomic divergence in sympatric and allopatric speciation of three Mihoutao (Actinidia) species. Hortic Res. 2022;9: uhac054. doi:10.1093/hr/uhac054

46. Irwin DE, Milá B, Toews DPL, Brelsford A, Kenyon HL, Porter AN, et al. A comparison of genomic islands of differentiation across three young avian species pairs. Mol Ecol. 2018;27: 4839–4855. doi:10.1111/mec.14858

47. Iwanaga S, Kaneko I, Kato T, Yuda M. Identification of an AP2-family Protein That Is Critical for Malaria Liver Stage Development. PLOS ONE. 2012;7: e47557. doi:10.1371/journal.pone.0047557

48. Vera IM, Beatty WL, Sinnis P, Kim K. Plasmodium Protease ROM1 Is Important for Proper Formation of the Parasitophorous Vacuole. PLOS Pathog. 2011;7: e1002197. doi:10.1371/journal.ppat.1002197

49. Al-Nihmi FMA, Kolli SK, Reddy SR, Mastan BS, Togiri J, Maruthi M, et al. A Novel and Conserved Plasmodium Sporozoite Membrane Protein SPELD is Required for Maturation of Exo-erythrocytic Forms. Sci Rep. 2017;7: 40407. doi:10.1038/srep40407

50. Real E, Nardella F, Scherf A, Mancio-Silva L. Repurposing of Plasmodium falciparum var genes beyond the blood stage. Curr Opin Microbiol. 2022;70: 102207. doi:10.1016/j.mib.2022.102207

51. van Schaijk BCL, Janse CJ, van Gemert G-J, van Dijk MR, Gego A, Franetich J-F, et al. Gene Disruption of Plasmodium falciparum p52 Results in Attenuation of Malaria Liver Stage Development in Cultured Primary Human Hepatocytes. PLOS ONE. 2008;3: e3549. doi:10.1371/journal.pone.0003549

52. Manzoni G, Marinach C, Topçu S, Briquet S, Grand M, Tolle M, et al. Plasmodium P36 determines host cell receptor usage during sporozoite invasion. Levashina E, editor. eLife. 2017;6: e25903. doi:10.7554/eLife.25903

53. Labaied M, Harupa A, Dumpit RF, Coppens I, Mikolajczak SA, Kappe SH. Plasmodium yoelii sporozoites with simultaneous deletion of P52 and P36 are completely attenuated and confer sterile immunity against infection. Infect Immun. 2007;75: 3758–3768.

54. Yuda M, Iwanaga S, Shigenobu S, Mair GR, Janse CJ, Waters AP, et al. Identification of a transcription factor in the mosquito-invasive stage of malaria parasites. Mol Microbiol. 2009;71: 1402–1414. doi:10.1111/j.1365-2958.2009.06609.x

55. Sasaki H, Sekiguchi H, Sugiyama M, Ikadai H. Plasmodium berghei Cap93, a novel oocyst capsule-associated protein, plays a role in sporozoite development. Parasit Vectors. 2017;10: 399. doi:10.1186/s13071-017-2337-8

56. Combe A, Moreira C, Ackerman S, Thiberge S, Templeton TJ, Ménard R. TREP, a novel protein necessary for gliding motility of the malaria sporozoite. Int J Parasitol. 2009;39: 489–496. doi:10.1016/j.ijpara.2008.10.004

57. Bosch J, Paige MH, Vaidya AB, Bergman LW, Hol WGJ. Crystal structure of GAP50, the anchor of the invasion machinery in the inner membrane complex of Plasmodium falciparum. J Struct Biol. 2012;178: 61–73. doi:10.1016/j.jsb.2012.02.009

58. Doi Y, Shinzawa N, Fukumoto S, Okano H, Kanuka H. ADF2 is required for transformation of the ookinete and sporozoite in malaria parasite development. Biochem Biophys Res Commun. 2010;397: 668–672. doi:10.1016/j.bbrc.2010.05.155

59. Kariu T, Yuda M, Yano K, Chinzei Y. MAEBL Is Essential for Malarial Sporozoite Infection of the Mosquito Salivary Gland. J Exp Med. 2002;195: 1317–1323. doi:10.1084/jem.20011876

60. Yang ASP, Lopaticki S, O’Neill MT, Erickson SM, Douglas DN, Kneteman NM, et al. AMA1 and MAEBL are important for Plasmodium falciparum sporozoite infection of the liver. Cell Microbiol. 2017;19: e12745. doi:10.1111/cmi.12745

61. Kumar V, Behl A, Sharma R, Sharma A, Hora R. Plasmodium helical interspersed subtelomeric family—an enigmatic piece of the Plasmodium biology puzzle. Parasitol Res. 2019;118: 2753–2766. doi:10.1007/s00436-019-06420-9

62. Warncke JD, Vakonakis I, Beck H-P. Plasmodium helical interspersed subtelomeric (PHIST) proteins, at the center of host cell remodeling. Microbiol Mol Biol Rev. 2016;80: 905–927.

63. Nkemngo FN, Raissa LW, Nguete DN, Ndo C, Fru-Cho J, Njiokou F, et al. Geographical emergence of sulfadoxine-pyrimethamine drug resistance-associated P. falciparum and P. malariae alleles in co-existing Anopheles mosquito and asymptomatic human populations across Cameroon. Antimicrob Agents Chemother. 2023;67: e00588–23.

64. Happi CT, Gbotosho GO, Folarin OA, Akinboye DO, Yusuf BO, Ebong OO, et al. Polymorphisms in Plasmodium falciparum dhfr and dhps genes and age related in vivo sulfadoxine–pyrimethamine resistance in malaria-infected patients from Nigeria. Acta Trop. 2005;95: 183–193.

65. Lefebvre MJM, Daron J, Legrand E, Fontaine MC, Rougeron V, Prugnolle F. Population Genomic Evidence of Adaptive Response during the Invasion History of Plasmodium falciparum in the Americas. Mol Biol Evol. 2023;40: msad082. doi:10.1093/molbev/msad082

66. Hamid MMA, Abdelraheem MH, Acheampong DO, Ahouidi A, Ali M, Almagro-Garcia J, et al. Pf7: an open dataset of Plasmodium falciparum genome variation in 20,000 worldwide samples. Wellcome Open Res. 2023;8: 22.

67. Dinerstein E, Olson D, Joshi A, Vynne C, Burgess ND, Wikramanayake E, et al. An Ecoregion-Based Approach to Protecting Half the Terrestrial Realm. BioScience. 2017;67: 534–545. doi:10.1093/biosci/bix014

68. Aleman JC, Fayolle A, Favier C, Staver AC, Dexter KG, Ryan CM, et al. Floristic evidence for alternative biome states in tropical Africa. Proc Natl Acad Sci. 2020;117: 28183–28190. doi:10.1073/pnas.2011515117

69. Rodríguez-Veiga P, Carreiras JMB, Quegan S, Heiskanen J, Pellikka P, Adhikari H, et al. Loss of tropical moist broadleaf forest has turned Africa’s forests from a carbon sink into a source. Sci Rep. 2025;15: 41744. doi:10.1038/s41598-025-27462-3

70. Nkemngo FN, Raissa LW, Nguete DN, Ndo C, Fru-Cho J, Njiokou F, et al. Geographical emergence of sulfadoxine-pyrimethamine drug resistance-associated P. falciparum and P. malariae alleles in co-existing Anopheles mosquito and asymptomatic human populations across Cameroon. Antimicrob Agents Chemother. 2023;67: e00588–23.

